# Orthosteric STING inhibition elucidates molecular correction of SAVI STING

**DOI:** 10.1101/2024.06.18.596881

**Authors:** Tao Xie, Max Ruzanov, David Critton, Leidy Merselis, Joseph Naglich, Jack Sack, Ping Zhang, Chunshan Xie, Jeffrey Tredup, Laurel B. Stine, Cameron Messier, Janet Caceres-Cortes, Luciano Mueller, Alaric J. Dyckman, John A. Newitt, Stephen C. Wilson

## Abstract

STING is broadly implicated in diseases ranging from cancer, autoimmune disease, neurodegeneration to rare, monogenic diseases.^1^ Early drug discovery campaigns focused on STING activation as a promising platform for cancer immunotherapy yet failed in multiple clinical trials due to lack of efficacy thus far.^2^ Current research and development activities concentrate on STING inhibition for treating autoimmune disease and neuroinflammation. While the progression of STING activators into the clinic has been successful, the discovery and clinical progression of STING inhibitors remain elusive. Questions persist about the molecular properties needed to distinguish between a STING activator and inhibitor, particularly within SAVI disease, a monogenic autoinflammatory disease that renders STING constitutively active.^3^ Here we leverage an orthosteric STING activator and inhibitor from the same chemical series to discover that STING M271 is a critical residue for molecular activation. The M271^CH3^ NMR chemical shifts reveal a unique molecular signature for pharmacological or genetically driven activation and inhibition that is not captured by x-ray crystallography. Additionally, M271 directly interacts with the most common SAVI mutation, V155M, and using an orthosteric STING inhibitor, we show partial rescue and molecular correction of STING V155M. Finally, these data present insights into therapeutic STING molecular correction for treating SAVI patients. Our results elucidate an unappreciated structural interaction critical for STING modulation that could be utilized as a molecular diagnostic tool for drug discovery. Furthermore, we demonstrate for the first time how the therapeutic requirements of a molecular corrector differ from an orthosteric STING inhibitor, and why this is important for the SAVI disease population.

---

The diABZI chemical family comprises molecules that can activate or inhibit STING signaling.^4,5^ To elucidate the differences in molecular architecture between orthosteric STING inhibition and activation, we determined the crystal structures of human STING^155-341^ bound to the orthosteric inhibitor, diABZI-i, and the orthosteric activator, diABZI-a1, derived from the diABZI series (Fig. 1a). Both compounds have similar potency for inhibiting and eliciting IFNβ in human PBMCs, respectively (IC50 = 49 ± 8 nM, 1 donor | EC50 = 117 ± 72 nM, 3 donors). However, the differences between the two structures are unexpectedly modest with a r.m.s.d. of 0.548 Å^2^ for all atoms (Fig. 1b, Supp. Fig. 1). Moreover, both compounds adopt similar positioning within the binding pocket with no obvious mechanistic rationale for their reciprocal functions evident in the crystal structures (Fig. 1c). Despite these findings, we analyzed both crystal structures for two characteristic structural features associated with STING activation: beta sheet lid formation and the apical wing distance between STING monomers (Fig. 1d).^6,7^

**Fig. 1.**
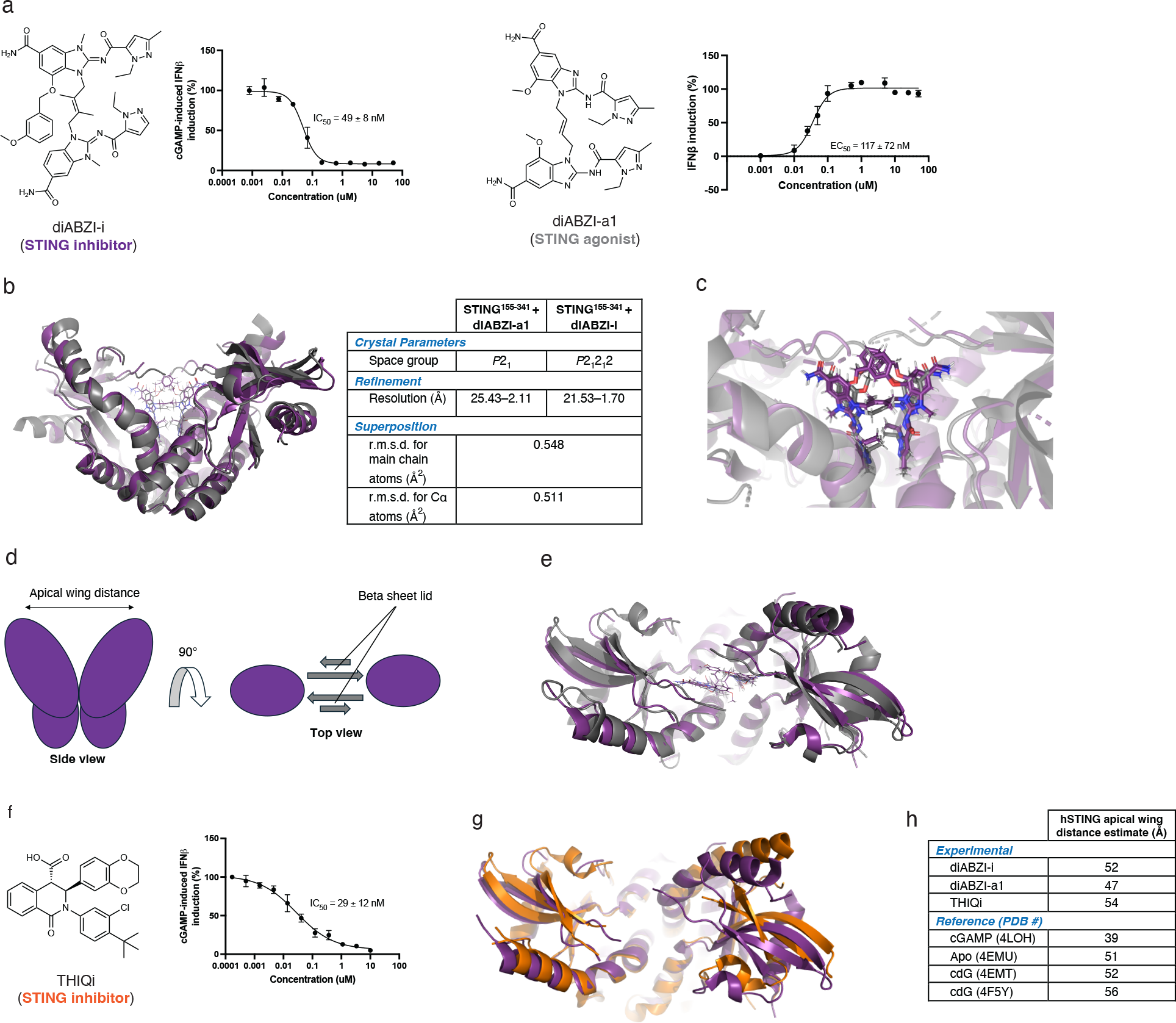
STING bound diABZI-based agonist and inhibitor are nearly indistinguishable in crystal structures. **(a)** diABZI-i inhibits cGAMP-induced IFNβ in PBMCs (n = 3 | 1 donor) and diABZI-a1 induces IFNβ in PBMCs (n = 3 | 3 donors, representative donor displayed). **(b)** Superposition of crystal structures (side view) of STING bound to diABZI-a1 (gray) and diABZI-i (purple). **(c)** diABZI-a1 and diABZI-i superposition over the STING binding pocket. **(d)** Cartoon depicting apical wing distance and beta sheet lid between STING monomers. **(e)** Superposition of crystal structures (top view) of STING bound to diABZI-a1 (gray) and diABZI-i (purple). **(f)** THIQi inhibits cGAMP-induced IFNβ in PBMCs (n = 3 | 1 donor). **(g)** Crystal structure overlay (top view) of STING bound to THIQi (orange) and diABZI-i (purple). **(h)** Apical wing distance measurements from structures in this study and reference structures collected from the PDB.

Beta sheet lid formation over the binding pocket is a well-described signature of cGAMP binding to STING.^6^ However, questions have been raised over whether its formation is necessary for driving STING signaling.^5^ Comparisons between STING bound to diABZI-i and diABZI-a1 do not reveal any significant differences in lid formation to explain inhibition or activation. Both compounds elicit a disordered STING lid organization that does not extend over the binding pocket (Fig. 1e), which has been previously described in both apo STING and STING bound to cyclic-di-GMP (cdG), a bacteria-derived STING agonist.^5,7–9^ Since orthosteric STING inhibition is poorly understood, we compared the STING bound between diABZI-i and THIQi, the only other orthosteric, chemically distinct STING inhibitor reported to date (Fig. 1f).^10^ Our findings show no significant differences in lid formation between these two structures (Fig. 1g), substantiating that beta sheet lid organization over the binding pocket is not required for orthosteric STING activation or inhibition.

Another well-reported feature of cGAMP-STING binding is the shortening of the apical wing distance between STING monomers.^6,7^ Both diABZI bound structures adopt a splayed open conformation with an apical wing distance of ∼47 Å and ∼52 Å for diAZBI-a1 and diABZI-i, respectively (Fig. 1h). Since the ∼5 Å difference between the two structures could be significant enough to predict STING inhibition or activation, we assessed these distances relative to reference structures of apo STING^8^, cGAMP-STING^6^, and cdG-STING^8,9^ in the PDB, and the THIQi-STING bound structure (Fig. 1h). All apical wing distances calculated in this study are bookended by cGAMP-STING (∼39 Å) and cdG-STING (∼56 Å), indicating that apical wing shortening or lengthening is not predictive for STING activation. Both orthosteric STING inhibitors used in this study adopt similar splayed open conformations, suggesting an open conformation could be required for inhibition. Therefore, the totality of our analysis indicates that STING^155–341^ crystallography cannot readily distinguish between diABZI-driven STING inhibition and activation.

Since small molecule binding to STING must drive conformational changes that lead to either signal transduction (agonist) or prevent signal transduction (inhibitor), we hypothesized that structural differences between diABZI-a1 and diABZI-i bound STING^155-341^ could be elucidated in the solution-state rather than solid-phase.^11^ To that end, we prepared ^13^C, ^15^N labelled STING^155-341^ for solution-state NMR spectroscopy. The THIQi-STING^155-341^ bound structure led to the highest quality spectra and therefore was used for ^1^H-^15^N HSQC (Supp. Fig. 2a) and ^1^H-^13^C HSQC (Fig. 2a) resonance assignments (see Methods section for detailed description of backbone and sidechain assignments, Supp. Fig. 2).

**Fig. 2.**
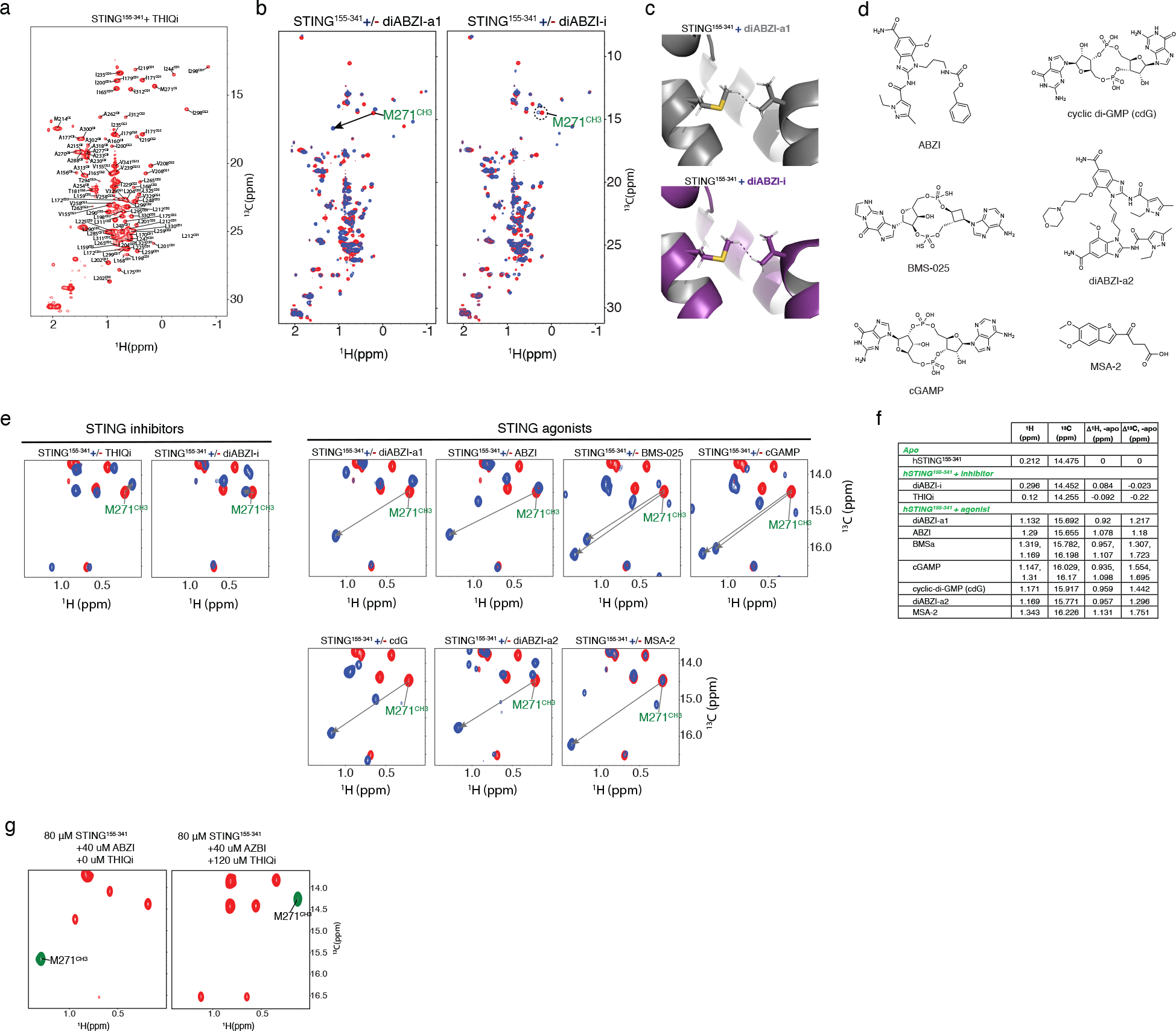
NMR distinguishes between STING activation and inhibition via M271^CH3^ chemical shift in ^1^H-^13^C HSQC spectrum. (a) ^1^H-^13^C HSQC sidechain methyl assignment of STING^155- 341^ + THIQi. **(b)** ^1^H-^13^C HSQC shifts of STING^155-341^ with and without diABZI-a1 and diABZI-i. **(c)** Crystal structures focused on V155 and M271 residues of STING^155-341^ bound with diABZI- a1 (upper, gray) and diABZI-i (lower, purple). **(d)** Chemical structures of diverse STING agonists used for this study. **(e)** Overlay of ^1^H-^13^C HSQC of STING^155-341^ recorded in the absence (red) and presence (blue) of various antagonists and agonists highlighting M271^CH3^. **(f)** Tabulated M271^CH3 1^H and ^13^C chemical shift differences between apo and bound STING. **(g)** ^1^H-^13^C HSQC competition experiment with ABZI (agonist) and THIQi (antagonist).

Comparison of ^1^H-^13^C HSQC spectra between diABZI-i and diABZI-a1 revealed multiple differences (Fig. 2b). Most notably, the methyl group on the M271 side chain (M271^CH3^) exhibited a significant downfield shift in both proton (0.920 ppm) and carbon (1.217 ppm) dimensions with diABZI-a1 but not diABZI-i. Re-examination of the diABZI-i and diABZI-a1 bound crystal structures exhibited no clear distinction between the two (Fig. 2c) with a r.m.s.d for the M271 residue <1 Å^2^ (Supp Fig 1a).

Major upfield shifts in both proton (0.721 ppm) and carbon (1.667 ppm) dimensions were also observed for A277^CH3^ upon binding to diABZI-a1 (Supp. Fig. 3) but not for diABZI-i, which is opposite in directionality to the M271^CH3^ shifts. Both M271^CH3^ and A277^CH3^ are >10 Å from the binding pocket as measured in the respective crystal structures. However, the magnitude and resolution of the M271^CH3^ shift compared to A277^CH3^ as well as its through-space proximity (2.5 Å) to V155 (Fig. 2c) caused us to focus our attention on M271^CH3^.

Since the differences in M271^CH3^ shifts between diABZI-i and diAZI-a1 were significant in magnitude and unobstructed by other correlations, we investigated whether this was a generalizable effect among other orthosteric STING inhibitors and agonists. We compared the STING-bound ^1^H-^13^C HSQC spectra among an additional seven chemically diverse STING agonists including cGAMP^12–15^, cdG^16^, ABZI^5^, diABZI-a2^5^, MSA-2^17^, and BMS-025^18,19^ (Fig. 2e) as well as THIQi. All STING agonists displayed significant downfield proton and carbon shifts (Fig. 2e, 2f, Supp Fig. 3). On average, STING agonists elicited a downfield proton shift of 1.016 ppm (min = 0.92 ppm, max = 1.131 ppm) and an average downfield carbon shift of 1.462 ppm (min = 1.18 ppm, max = 1.751 ppm). On the other hand, both STING inhibitors showed only modest effects on proton and carbon chemical shifts. STING inhibitors showed no appreciable effect on the proton chemical shift on average (-0.004 ppm) and a modest average upfield carbon shift of -0.122 ppm (Fig. 2e, 2f, Supp. Fig. 3). Incubation with a STING agonist followed by an excess of inhibitor elicited a downfield shift of M271^CH3^ followed by an upfield shift in the ^1^H-^13^C HSQC spectrum (Fig. 2g). This is consistent with a competitive rescue following STING activation and validates that the M271^CH3^ chemical shift can be diagnostic for STING activation and inhibition.

**Fig. 3.**
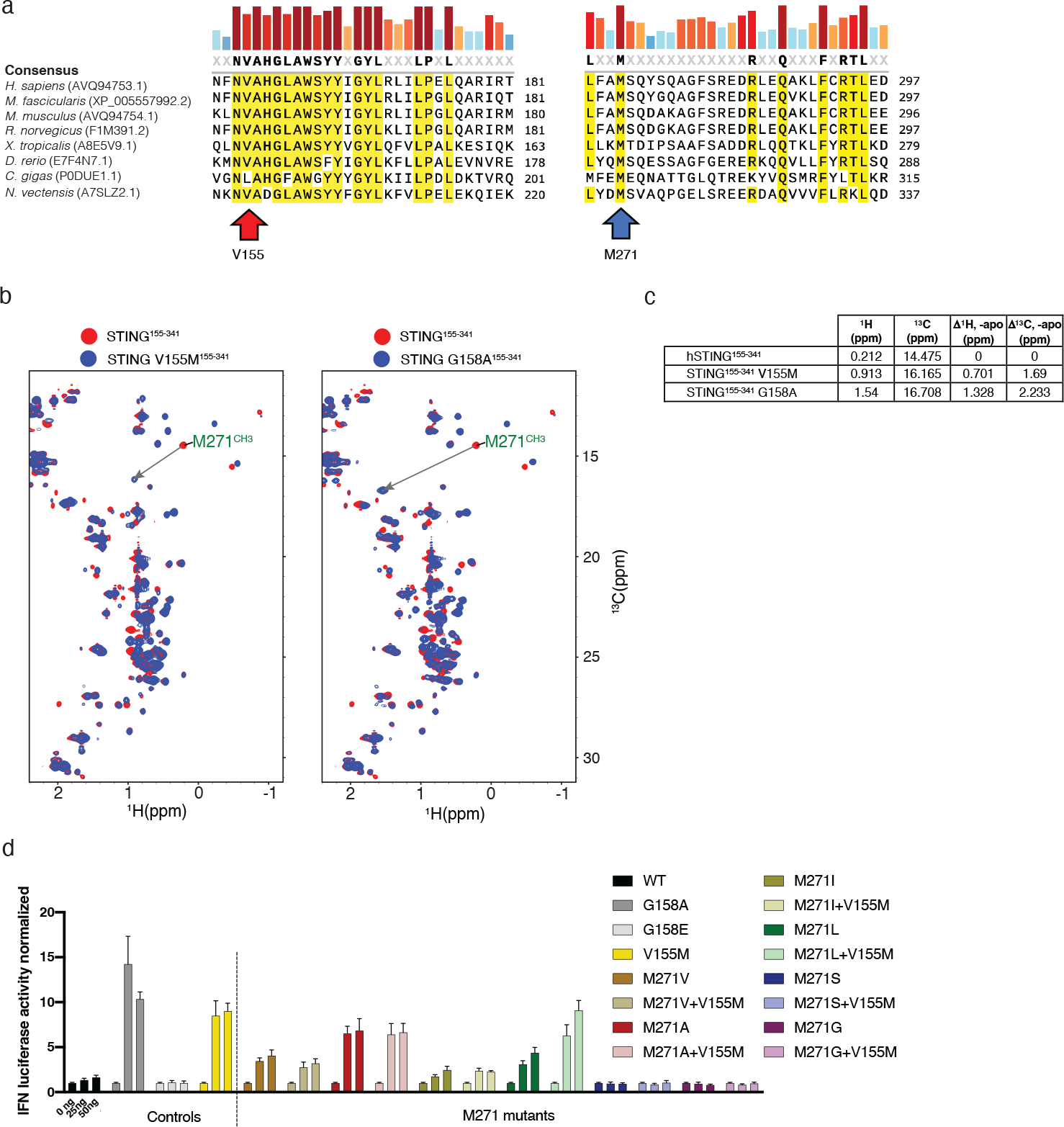
The evolutionary conserved M271 residue mediates SAVI STING V155M-driven IFN induction. (a) V155 and M271-focused phylogenetic sequence alignment. **(b, c)** Comparison of ^1^H-^13^C HSQC spectra among STING WT, SAVI STING V155M and SAVI STING G158A. **(d)** M271-focused mutational analysis using HEK293 IFN-driven luciferase reporter cell line (n = 3).

Since M271^CH3^ and A277^CH3^ are >10 Å from the binding pocket, it is likely that the large chemical shift movements are caused by agonist binding-induced conformational changes rather than direct interaction. By mapping the chemical shift differences between STING-bound ABZI and THIQi onto crystal structures, we found that the hydrophobic region harboring M271 and A277 beneath the CDN binding site is significantly perturbed and indicative of distinct conformations between agonist and antagonist-bound STING^155-341^ (Supp. Fig. 4a, b). To understand why agonist binding induced such large shifts but not antagonists, we compared crystal structures of apo-STING (PDB: 4EMU) and THIQi-bound STING with 2’,3’-cGAMP-bound STING. In the inactive state (i.e. apo and THIQi-bound STING), M271^CH3^ sits directly in between the aromatic rings of W161 and F279. Here it forms sulfur-aromatic interactions, which leads to a large shielding effect caused by the aromatic ring current and subsequent upfield shift. (Supp. Fig. 4c, d). A277^CH3^ resides above, off center the indole ring of W161 and proximal and parallel to the benzene ring of Y274 (Supp. Fig. 4c, d). In the active state (i.e. cGAMP-bound STING), the M271^CH3^ sulfur-aromatic interaction is disrupted due to a shift in relative orientation between M271^CH3^ and the aromatic rings of W161 and F279. M271^CH3^ experiences deshielding by moving outside the indole rings of W161 and F279 (Supp. Fig 4e), leading to downfield shift. In comparison, A277^CH3^ experiences large shielding by moving to the top center of the W161 indole ring and distal from the Y274 ring, resulting in upfield shifts. These local conformational differences between active and inactive state clearly explain the respective large downfield and upfield shifts of M271^CH3^ and A277^CH3^ upon binding to agonists and the small chemical shift changes upon antagonist binding.

The STING V155M mutation is reported to be the most common genotype driving STING-Associated Vasculopathy with onset in Infancy (SAVI), a severe autoimmune disease with poor treatment options available.^3,20^ Approximately 62% of known SAVI patients harbor the V155M mutation, and ∼86% of reported SAVI patients have mutations in the same region as V155M, highlighting a crucial area for eliciting STING activation.^20^ The through-space proximity of M271 to V155 (2.5 Å) suggests an important relationship between these two residues (Fig. 2d), and the phylogenetic conservation of M271 between invertebrates and vertebrates indicates a preservation of its importance over the evolution of STING signaling (Fig. 3a). Computational modeling also suggests an important interaction between V155M and M271 that reinforces STING dimer stability.^21^ Therefore, we sought to investigate whether the M271 NMR signature for pharmacologically induced STING activation is also observed in genetically induced STING activation through SAVI mutations.

We prepared ^13^C, ^15^N labelled STING^155-341^ V155M and G158A^22,23^ for solution-state NMR spectroscopy. Both SAVI mutations elicited significant M271 downfield chemical shift changes consistent with those illustrated with pharmacological activation (Fig. 3b). STING^155-341^ V155M elicited a downfield proton shift of 0.701 ppm and a downfield carbon shift of 1.69 ppm, whereas STING^155-341^ G158A caused a downfield proton shift of 1.328 ppm and a downfield carbon shift of 2.233 ppm (Fig. 3c). We also examined the NMR profile of STING^155-341^ G158E, a rationally designed mutation at a SAVI location that does not elicit constitutive STING signaling.^23^ However, the M271^CH3^ peak in the ^1^H-^13^C HSQC spectrum could not be identified (Supp Fig. 5), which may be due to STING destabilization from steric bulk or negative charge as previously suggested.^23^ This data shows that the M271^CH3^ chemical shift can be leveraged as a diagnostic signature for pharmacological or genetic STING activation.

The relationship between M271 and SAVI V155M is a provocative connection that we wanted to leverage to gain insight into SAVI for its potential treatment. As such, we decided to evaluate the consequence of M271 mutagenesis on constitutive STING interferon (IFN) signaling with and without a V155M background. M271 mutagenesis to alternative hydrophobic amino acids, M271A, M271I, M271L, and M271V, conferred constitutive activation to STING on par with V155M-driven signaling (Fig. 3d). Our functional assay indicates that M271 critically controls signaling induced by the V155M mutation. M271 mutagenesis to M271A, M271I and M271V controlled the amplitude of V155M induced signaling. M271L is distinctive in that its activity under a V155M background is enhanced compared to M271 alone, yet M271L + V155M is attenuated compared to V155M alone. Furthermore, M271S and M271G completely ablated V155M-driven signaling, confirming that M271 is critical residue for V155M SAVI signal transduction (Fig. 3d).

M271 mutagenesis can confer a gain-of-function state suggesting that mutation of this residue could confer SAVI-like disease. While no clinical publication has been described to date on such a patient population, we identified one heterozygous carrier of M271I within the gnomAD database^24^. It is currently unknown whether this carrier presents with SAVI-like phenotypes, but it is notable to mention that Polyphen^25^ and SIFT^26^ annotate M271I as a “possibly damaging” and “deleterious” mutation, respectively.

Given that the M271^CH3^ chemical shifts act as a diagnostic for STING activation, we decided to examine whether orthosteric STING inhibition could molecularly correct the M271^CH3^ activation shift of V155M SAVI.^27,28^ Both diABZI-i and THIQi showed partial upfield corrections of the STING V155M M271^CH3^ chemical shift (Fig. 4a, 4b). diABZI-i exhibited a Δ^1^H = -0.284 ppm, Δ^13^C = -0.748 ppm correction, and THIQi led to a Δ^1^H = -0.47 ppm, Δ^13^C = - 0.514 ppm correction (Fig. 4c). All STING agonists led to further downfield proton and carbon shifts of V155M M271^CH3^ (Fig. 4c) except for ABZI, where only a downfield proton shift was apparent with no significant change in the carbon chemical shift.

**Fig. 4.**
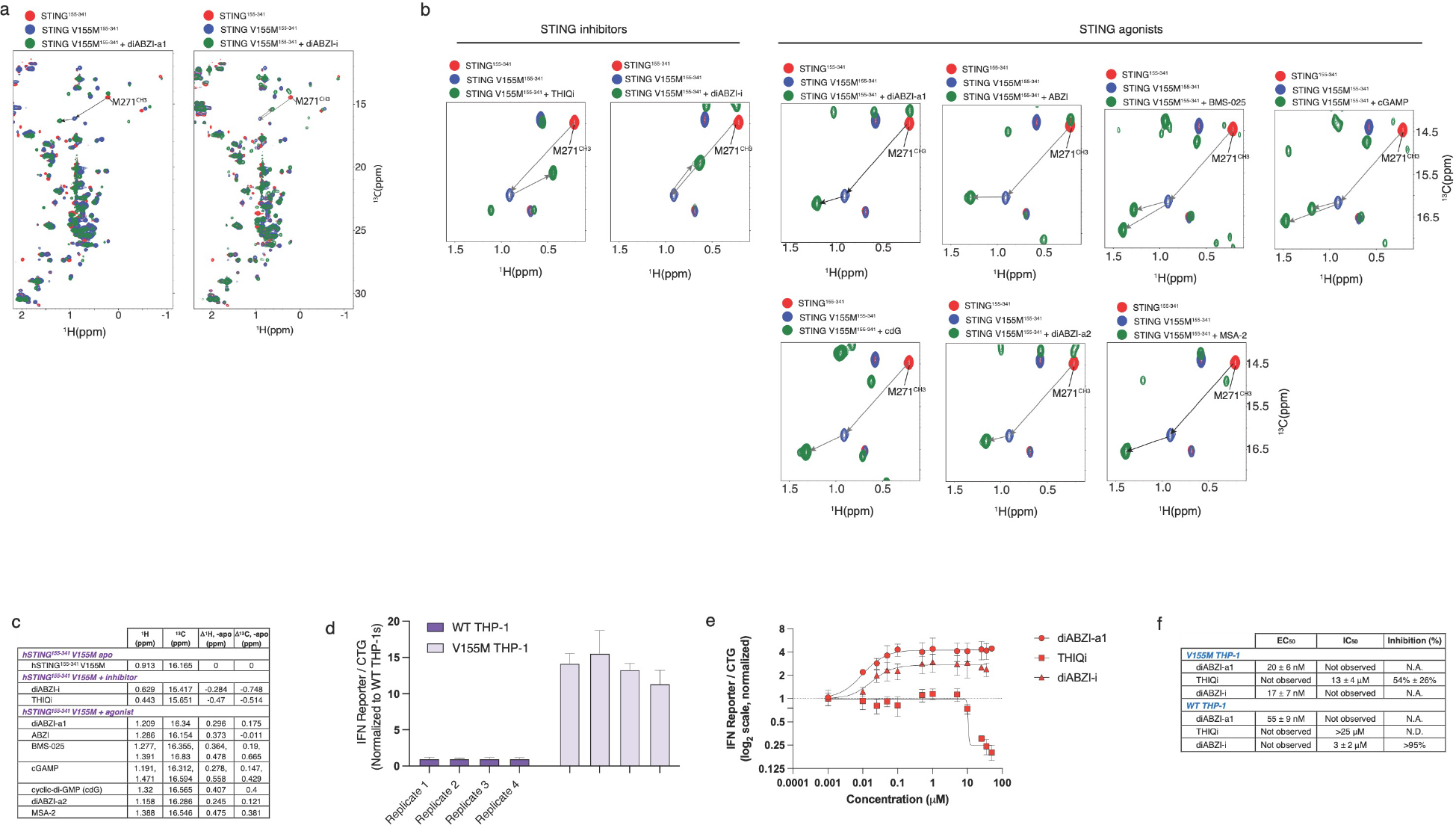
M271^CH3^ chemical shift correction is necessary but not sufficient for STING molecular correction of V155M SAVI. (a + b) Overlay of ^1^H-^13^C HSQC of STING^155-341^ V155M recorded in the absence (red) and presence (blue) of various antagonists and agonists. **(c)** Tabulated M271^CH3 1^H and ^13^C chemical shift differences between apo and bound STING V155M. **(d)** Comparison of basal IFN-driven luciferase levels between WT THP-1 and STING V155M THP-1 cell lines **(e)** Agonism and antagonism behavior of diABZI-a1, THIQi, and diABZI-i on basal IFN levels in STING V155M THP-1 cell line (n = 3); representative replicate shown. **(f)** Tabulated IFN-driven luciferase EC50 and IC50 values in WT THP-1 (with and without cGAMP stimulation) and STING V155M THP-1 (no stimulation).

SAVI patient carriers present with symptoms consistent with a type I interferonopathy. The disease often manifests in early life leading to systemic inflammation and a high mortality rate with features including elevated type I IFN, skin vasculopathy, arthritis, and pulmonary disease.^3,20^ The pulmonary aspect of SAVI disease is the most common feature among patients and a key driver for mortality. In published studies to date, JAK inhibitors such as ruxolitinib are the most reported treatment for SAVI.^3,29–31^ However, it is important to note that while treatment with ruxolitinib or other JAK inhibitors is reported to improve clinical disease scores, reports indicate little to no impact on clinical IFN scores.^20,29,31^ While more work is need to understand this paradox, it implies that targeting STING could be the only critical node for therapeutically resolving SAVI disease.

Since THIQi and diABZI-i show partial molecular correction of SAVI V155M STING by M271 in the ^1^H-^13^C HSQC spectrum, we were interested in understanding whether orthosteric STING inhibition could also reverse V155M SAVI constitutive activation in a cellular model. A stably expressed STING V155M THP-1 model exhibits elevated IFN (>10x) over WT THP-1 in the absence of cGAMP stimulation (Fig. 4d). THIQi demonstrates partial rescue of the elevated IFN in this model (IC50 = 13 ± 4 μM, % inhibition = 54% ± 26%) with an inhibition potency within three-fold of the WT THP-1 cGAMP stimulation model (Fig. 4e, 4f). In contrast, diABZI- i demonstrates expected inhibition in the WT THP-1 cGAMP stimulation model (IC50 = 3 ± 2 μM) but potent agonism in the V155M SAVI model (EC50 = 17 ± 7 nM) with a >170x increase in potency relative to inhibition potency in WT THP-1s. These data suggest orthosteric STING inhibition may not necessarily resolve the constitutive signaling associated with V155M SAVI, and in fact cautions that an orthosteric STING inhibitor in WT carriers could serve as a STING agonist in SAVI carriers, thereby exacerbating disease.

Cryo-EM studies of full-length STING have examined the important interplay between the cyclic dinucleotide (CDN) binding region and transmembrane region of STING, leading to oligomerization-dependent effects on signaling.^22,23,32–34^ However, STING has multiple signaling mechanisms embedded and distributed throughout its architecture, which makes therapeutic inhibition challenging.^35–38^ The chemical shift of M271^CH3^ elucidates STING signaling and provides a robust diagnostic for STING activation, inhibition, and molecular correction conferred by both genetics and pharmacology. In the case of V155M SAVI, this has exposed the challenges and potential risks with treating V155M SAVI patients using orthosteric STING inhibitors. In fact, orthosteric STING targeting of V155M SAVI requires molecular correction not just inhibition.

## METHODS

### Frozen human PBMC cytokine production assays

Frozen human peripheral blood mononuclear cells (PBMCs, BioIVT) are gently thawed, suspended in complete medium [AIMV, 10% heat-inactivated FBS, 1% antibiotic-antimycotic, 1% sodium pyruvate, 1% MEM non-essential amino acids; ThermoFisher] and concentrated by low-speed centrifugation (400 x g, ambient temperature, five minutes). The cell pellet is suspended in complete medium and incubated for one hour (37 °C, 5% CO2) prior to measuring the cell concentration and viability. For each assay, compounds, suspended in DMSO, are first transferred to empty wells of a 384-well plate, followed by adding PBMCs suspended in complete medium to each well (100,000 cells / well for IFN-β assays). Cells are pre-treated for one hour (37 °C, 5% CO2). After one-hour, complete medium is added to baseline control wells or 2′3′-cGAMP (100 μM, Chemietek) is added to designated 100% maximum control and test compound wells. Cells are subsequently treated for 24 hours prior to transferring the supernatants to empty 384-well white, solid-bottom assay plate wells. Cellular interferon-beta was then measured using alphalisa immunoassays (Revvity) following vendor protocols. The half maximal inhibitory concentration values (IC50; compound concentration which inhibits a response halfway between the assay baseline and maximum) are calculated using PRISM software.

### Fresh human PBMC cytokine production assays

Peripheral blood was collected from healthy volunteers into sodium heparinized vacutainer CPT tubes (BD Biosciences) and shaken at room temperature until centrifugation. CPT tubes were centrifuged at 1800 x g for 20 minutes at 23 °C resulting in the separation of PBMCs from granulocytes and red blood cells. The PBMC layer was carefully transferred into a 15 mL conical tube and PBS was added to bring the total volume to 15 mL, then inverted to mix, and centrifuged at 300 x g for 15 minutes. Supernatant was aspirated without disturbing the cell pellet, then cells were resuspended in 10 mL PBS and centrifuged at 300 x g for 10 minutes.

Supernatant was aspirated and cells were resuspended in complete medium consisting of RPMI 1640, 10% heat-inactivated FBS, and 100 U/mL Pen-Strep, transferred to T25 flasks, and rested at 37 °C with 5% CO2 overnight. The following day, cells were transferred to a 15 mL conical tube, counted, and adjusted to 1E+06 cells/mL in complete medium. 100 uL of cell suspension (1E+05 cells) was dispensed into each well of a U-bottom 96-well plate. Next, 10 mM stock compounds dissolved in DMSO were dispensed using a D300e Digital Dispenser (HP) to generate 11-point dose-response curves ranging from 1.3 nM to 50 μM, with DMSO-only representing the baseline; each point was analyzed in duplicate. All wells were normalized with DMSO equivalent to the 50 μM concentration volume and total DMSO was kept below 1% of the final volume. Plates were then incubated at 37°C with 5% CO2 for 24 hours. Supernatant was collected and used for interferon-beta detection by AlphaLISA (Revvity) following manufacturer’s instructions, and cell viability was assessed by CellTitre-Glo assay (Promega).

### STING expression and purification

Human STING^155−341^ G230A/R293Q, STING^155−341^ V155M/G230A/R293Q, STING^155−341^ G158A/G230A/R293Q, and STING^155−341^ G158E/G230A/R293Q were subcloned into a pET28 bacterial expression vector with a construction containing an N-terminal hexahistidine tag followed by a SUMO tag fusion and a Tobacco Etch Virus (TEV) protease cleavage site. Protein expression and purification were carried out essentially as described previously^9^. In brief, the STING expression vector was transformed into chemically competent *E. coli* One Shot BL21 Star (DE3) cells (Invitrogen). The unlabeled expression was carried out in the Terrific Broth medium (Gibco). The ^13^C, ^15^N labeled protein was expressed in a modified M9 medium containing D-glucose (Cambridge Isotope Laboratories, Inc., U-^13^C6, 99%) and ammonium chloride (Cambridge Isotope Laboratories, Inc., ¹⁵N, 99%). The cells were grown in shake-flasks at 37 °C to an OD^600 nm^ of 1 AU, and the expression was induced with 0.4 mM isopropyl β-D-1- thiogalactopyranoside. The induced cells were harvested after overnight incubation at 18 °C. The cells were lysed, crude protein was captured using a HisTrap FF Crude column (Cytiva), and the eluted protein was further purified using a HiLoad 26/600 Superdex 200 gel filtration column (Cytiva). Fractions from the single, major peak fractions containing enriched STING protein were pooled, and incubated with TEV protease overnight cleavage at 4°C to cleave off the His-SUMO fusion. The untagged STING was isolated in the flow through following chromatography over Ni-NTA (Qiagen) resin. The final, purified STING protein was buffer exchanged into storage buffer (25 mM Tris 7.5, 200 mM NaCl, 0.5 mM TECP, 5% glycerol) using a PD10 column (Cytiva). Purified STING in storage buffer was flash-frozen in liquid nitrogen and stored at −80 °C until needed.

### X-ray crystallographic data collection, processing, and structure determination of STING co-crystallized with diABZI-a1 and diABZI-i

Purified hSTING^155-341^ G230A/R293Q was co-crystallized with diABZI-a1 as follows. STING (at a protein concentration of 10 mg/mL in 20 mM HEPES pH 7.5, 300 mM NaCl, 1 mM TCEP) was combined with diABZI-a1 (final ligand concentration = 1.5 mM), and the mixture incubated on ice for 1 hour. Crystallization screening was carried out by combining 0.2 µL STING + diABZI-a1 mixture with an equal volume of precipitant (0.2 M ammonium acetate, 0.1 M bis-tris pH 5.5, 25% (w/v) PEG 3,350) over a reservoir of 75 µL precipitant in an MRC UVXPO sitting drop vapour diffusion crystallization tray (Swissci) at room temperature. Purified hSTING^155-341^ G230A/R293Q was co-crystallized with diABZI-i as follows. STING (at a protein concentration of 10 mg/mL in 20 mM HEPES pH 7.5, 300 mM NaCl, 1 mM TCEP) was combined with diABZI- i (at a final ligand concentration of 1.5 mM), and the mixture incubated on ice for 1 hour. Crystallization screening was carried out by combining 0.2 µL STING + diABZI-i mixture with an equal volume of precipitant (0.2 M magnesium chloride hexahydrate, 0.1 M bis-tris pH 5.86, 25% (w/v) PEG 3,350) over a reservoir of 75 µL precipitant in an MRC UVXPO sitting drop vapour diffusion crystallization tray (Swissci) at 4 °C. Crystals typically formed within 24 hours and grew to maximum size within 1 week. Crystals were harvested using a nylon loop, cryoprotected in precipitant supplemented with 25% (v/v) glycerol and were then flash frozen in liquid nitrogen for X-ray diffraction screening. X-ray diffraction data for STING + diABZI-a1 was collected at a wavelength of 0.92 Å and a temperature of 100 K at the National Synchrotron Light Source II (AMX beamline 17-ID-1). X-ray diffraction data for STING + diABZI-i was collected at a wavelength of 1 Å and a temperature of 100 K at the Advanced Photon Source (IMCA-CAT beamline 17-ID). Data reduction was performed using autoPROC (Global Phasing Ltd.). Structures were phased by molecular replacement using Phaser^39^ and the protein coordinates from a BMS-internal crystal structure of STING determined previously. The structure was completed through iterative cycles of model building using Coot^40^ and restrained refinement using autoBUSTER (Global Phasing Ltd.). A summary of the data collection and refinement statistics is provided in Table 1. The crystal structures of STING with diABZI-a1 and diABZI-i have been deposited into the RCSB PDB under accession numbers **####** and **####**, respectively.

**Table 1.**
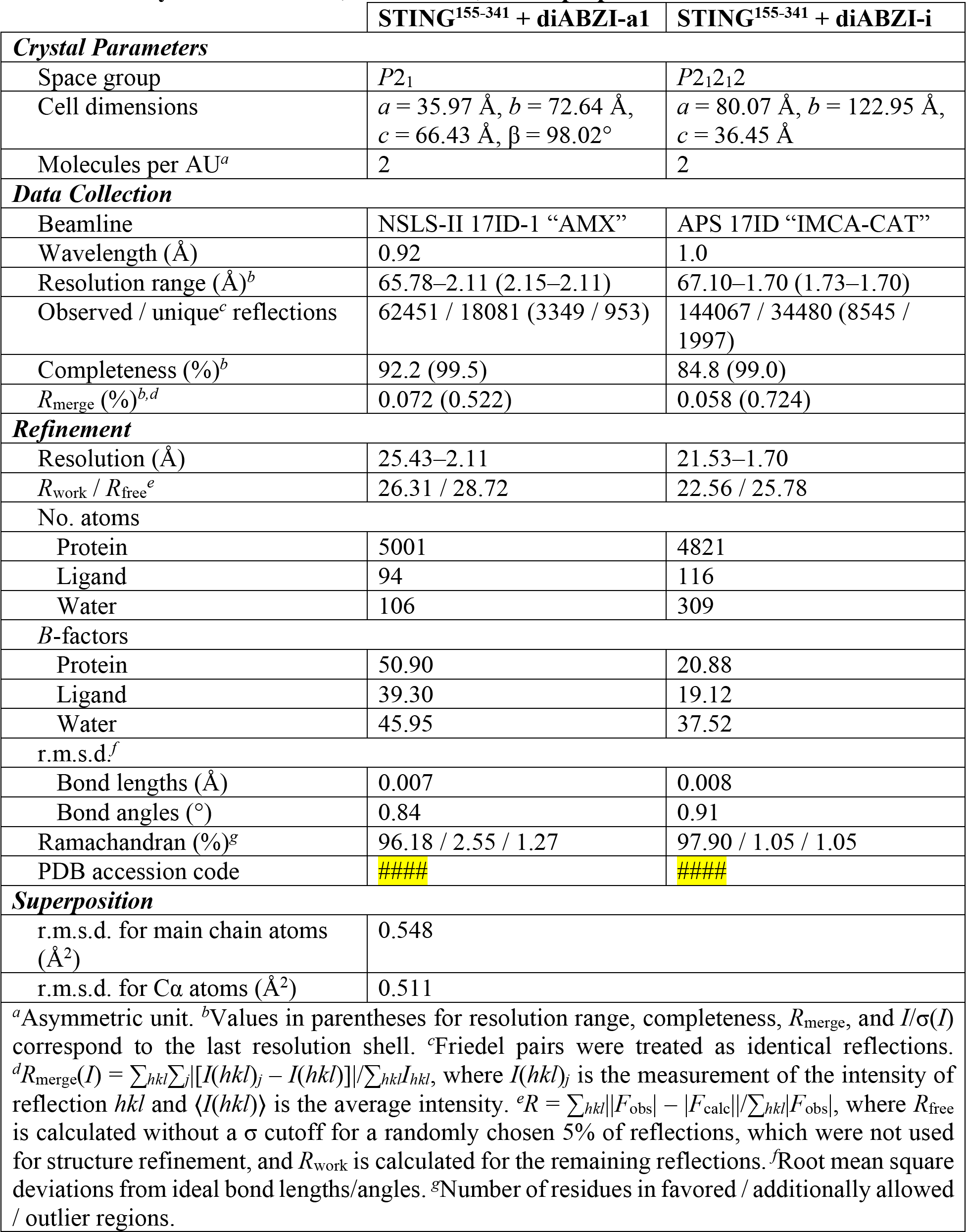
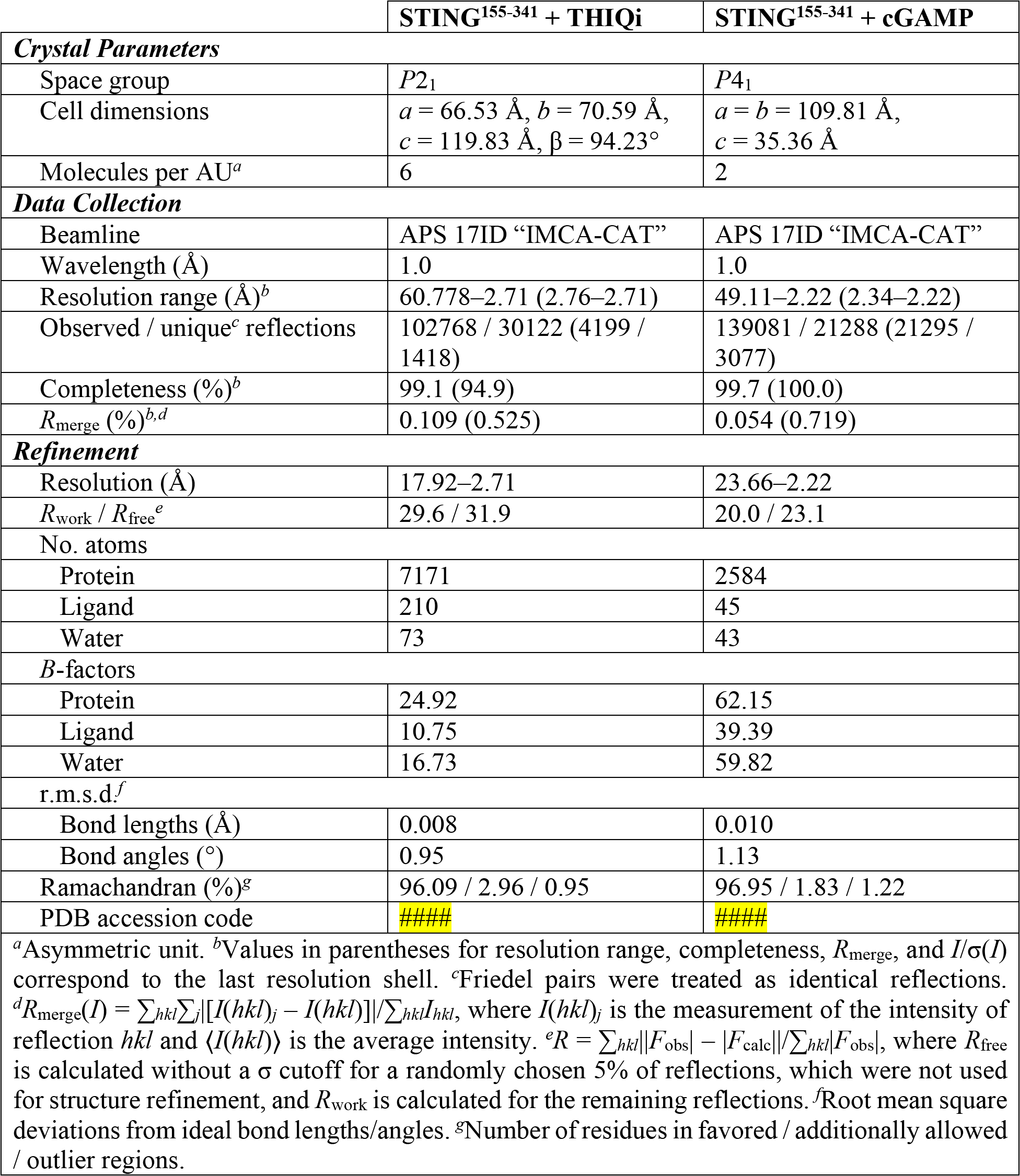
X-Ray Data Collection, Refinement, and Superposition Statistics.

### NMR sample preparation

The 0.8 mM STING^155-341^ G230A/R293Q -THIQi complex for backbone and methyl assignments was generated by mixing ^13^C, ^15^N-labeled STING^155-341^ G230A/R293Q and THIQi (1:6) at ∼10 µM concentration in 20 mM Tris-d11 (pH 7.3) including 100 mM NaCl, 3mM dithiothreitol-d10, 8% D2O, followed by ultrafiltration to remove extra compound. The ^13^C, ^15^N- labeled STING^155-341^ G230A/R293Q -ABZI complex was made in the same way as STING^155-341^ G230A/R293Q -THIQi complex, and the final concentration is ∼0.95 mM. ^2^H, ^13^C,^15^N labeled STING^155-341^ G230A/R293Q was unfolded in 20 mM Tris-d11 (pH 7.3) including 100 mM NaCl, 8 M Urea, 3 mM dithiothreitol and followed by refolding in the presence of ABZI in the same buffer except without urea by dilution at 4 °C. The ^2^H, ^13^C,^15^N labeled STING^155-341^ G230A/R293Q -ABZI was used for backbone assignment. Samples of the complexes were buffer exchanged to 99.96% D2O for experiments in D2O.

### NMR spectroscopy and assignments

NMR experiments for assignment of THIQi and ABZI-complexed STING^155-341^ G230A/R293Q were acquired at 35 °C and 30 °C respectively on Bruker Ultrashield^TM^ 700 MHz and Ascend^TM^ 600 MHz spectrometers. NMR data processing and analysis were performed using NMRPipe^41^ and NMRFAM-Sparky^42^. Backbone ^1^H, ^13^C, and ^15^N and sidechain methyl assignments of STING^155-341^ G230A/R293Q -THIQi complex were obtained by analyzing 2D ^1^H-^15^N TROSY and ^1^H-^13^C constant-time HSQC (CT-HSQC) and three-dimensional HNCA, HN(CO)CA, HNCO, HN(CA)CO, ^15^N-edited NOESY-HSQC, and ^13^C-edited NOESY-HSQC spectra collected on ^15^N,^13^C labeled sample. Backbone and methyl resonance assignments of STING^155-341^-ABZI complex were achieved by analyzing 2D ^1^H-^15^N TROSY and 3D HNCACB, HNCA, HN(CO)CA, HNCO, and HN(CA)CO spectra collected on ^2^H, ^13^C,^15^N labeled sample plus 2D ^1^H-^15^N TROSY, ^1^H-^13^C CT-HSQC and 3D ^15^N-edited NOESY-HSQC, ^13^C-edited NOESY-HSQC spectra acquired on ^13^C,^15^N labeled sample.

In ^1^H-^13^C CT-HSQC experiment, methyls with an odd number aliphatic carbon neighbors show opposite intensity compared to those with an even number aliphatic carbon neighbors.^43,44^ Since the CH3 group of Met has no aliphatic carbon neighbors, its NMR signal intensity is opposite to those of Ala, Ile, Leu, Thr, and Val in methyl fingerprint region of NMR spectrum.

Therefore, chemical shifts of Met methyls are easy to be identified. Assisted with 3D ^13^C-edited NOESY-HSQC, M271^CH3^ and A277^CH3^ in both complexes were unambiguously assigned (Fig. 2a, Supp. Fig. 2b, 2d and 2e). The backbone and methyl assignments of STING^155-341^ G230A/R293Q in complex with other ligands were achieved by comparing with 2D ^1^H-^15^N TROSY and ^1^H-^13^C HSQC of these two complexes.

### NMR titrations

50-100 µM of ^13^C, ^15^N-labeled STING^155-341^ G230A/R293Q, STING^155−341^ V155M/G230A/R293Q, STING^155−341^ G158A/G230A/R293Q, and STING^155−341^ G158E/G230A/R293Q was titrated with different ligands in 20 mM Tris-d11 (pH 7.3) including 100 mM NaCl, 3 mM dithiothreitol-d10, 8% D2O, and the 2D ^1^H-^15^N TROSY and ^1^H-^13^C HSQC experiments were collected at 25 °C.

### HEK cell culture and luciferase reporter assay

HEK-Lucia^TM^ Null Cells were obtained from Invivogen. HEK-Lucia^TM^ cells were maintained in Dulbecco’s modified Eagle’s medium (DMEM) supplemented with 10% heat-inactivated fetal bovine serum (Thermo Fisher Scientific), penicillin and Normocin^TM^ (100 U/mL and 100 ug/mL). HEK-Lucia^TM^ cells were transfected with STING wild type, or mutant constructs using Lipofectamine^TM^ 3000 (Thermo Fisher Scientific). After a 20h incubation, luciferase activity was determined using QUANTI-Luc^TM^ following the standard protocol (Invivogen).

### THP-1 cell culture and luciferase reporter assay

THP1-Dual^TM^ KI STING cells harboring either the WT allele or V155M point mutation were acquired from Invivogen (cat# thpd-m155). Both cell lines were maintained and passaged according to manufacturer’s recommendation, briefly, cells were grown in RPMI 1640 supplemented with 10% heat-inactivated FBS, and Pen-Strep (100 μg/mL). Following recovery from cryopreservation (>2 passages), cells were sub-cultured and passaged every 3 days.

Selection pressure was maintained by addition of 10 μg/mL blasticidin and 100 μg/mL zeocin. All experiments were conducted on cells under 15 passages to maintain cell line stability.

Cells were seeded in 96 well plates at a density of 100,000 cells in 100 μL of media. When specified stimulation was conducted by overlaying 100 μM of 2′3′ cGAMP (cat# tlrl- nacga23 dissolved in LAL water) on cells for 24 hours following 1 hour pre-incubation with STING inhibitor or agonist. Following incubation, plates were spun down at 500 x g for 5 mins, 20 μL of cell supernatant was collected and added to white opaque plates. 50 μL of freshly prepared QUANTI-Luc reagent 4 (cat# rep-qlc4lg1) was added to the cell supernatant and end point luminescence was read immediately on an Envision plate reader. Cell viability for each study was assessed by treating cells with CellTiter-Glo (cat# G9242) according to manufacturer’s protocol and luminesce was quantified.

### THP1-Dual^TM^ cell reporter assays

Engineered human THP1-Dual cells (Invivogen), derived from the human THP1 monocyte cell line, contain two stably integrated inducible reporter genes measuring interferon (luciferase production) and NF-κB (secreted embryonic alkaline phosphatase production, SEAP) promoter activities. The cells are maintained following the supplier protocols in complete culture medium [RPMI 1640, 10% heat-inactivated fetal bovine serum (FBS), 1% penicillin/streptomycin 10 ug/ml blasticidin, 1 mM sodium pyruvate, 100 ug/ml zeocin; ThermoFisher].

*Antagonist-Mode*: Compounds suspended in DMSO or DMSO alone are first transferred to empty wells of a 384-well plate. Assay medium [RPMI 1640, 1% penicillin/streptomycin, 0.1% bovine serum albumin (Sigma)] is added to baseline control wells and cells suspended in assay medium are added to wells containing DMSO (100% maximum control) or test compound (15,000 per well). Cellular STING signaling is then induced by adding 2′3′-cGAMP (Chemietek), at final concentrations of 15 μM for interferon (luciferase) assays. After 24 h treatment (37°C, 5% CO2), cell viability is measured using CellTiter Fluor (Promega) and cellular luciferase is measured by adding freshly prepared Quanti-Luc (luciferase, Invivogen) following manufacturer’s instructions. The half maximal inhibitory concentration values (IC50; compound concentration which inhibits a response halfway between the assay baseline and maximum) are calculated using PRISM software.

*Agonist-Mode*: Compounds or DMSO alone are transferred to empty wells of a 384-well plate. Assay medium is added to baseline control wells and THP1-dual cells suspended in assay medium are added to wells containing DMSO alone (100% maximum control) or test compound (15,000 per well). Interferon-beta (400 IU/mL; R&D Systems) is used to establish 100% maximum values. After 24 h treatment (37 °C, 5% CO2), cell viability is measured using CellTiter Fluor (Promega) and cellular luciferase is measured using Quanti-Luc. The half maximal effective concentration values (EC50; compound concentration which elicits a response halfway between the assay baseline and maximum) are calculated using PRISM software.

### Compound dosing

When specified, STING modulating compounds were dosed using the Tecan D300e digital dispenser. Compound stocks were formulated in DMSO at 10 mM. DMSO normalization on non-treated wells was conducted by automatic dispenser.

## ACKNOWLEDGEMENTS

The authors thank Bill Degnen and Cynthia Hendrix for *in vitro* cellular assay support as well as our Cell Technologies and Compound Management Groups for routinely providing cell and compound supplies support. We thank Dr. Arvind Mathur for supporting the study. We also thank Mary Struthers for her insights into the paper as well as Sophie Roy for insightful conversations on the topic.

## EXTENDED FIGURES

**Extended Fig. 1.**
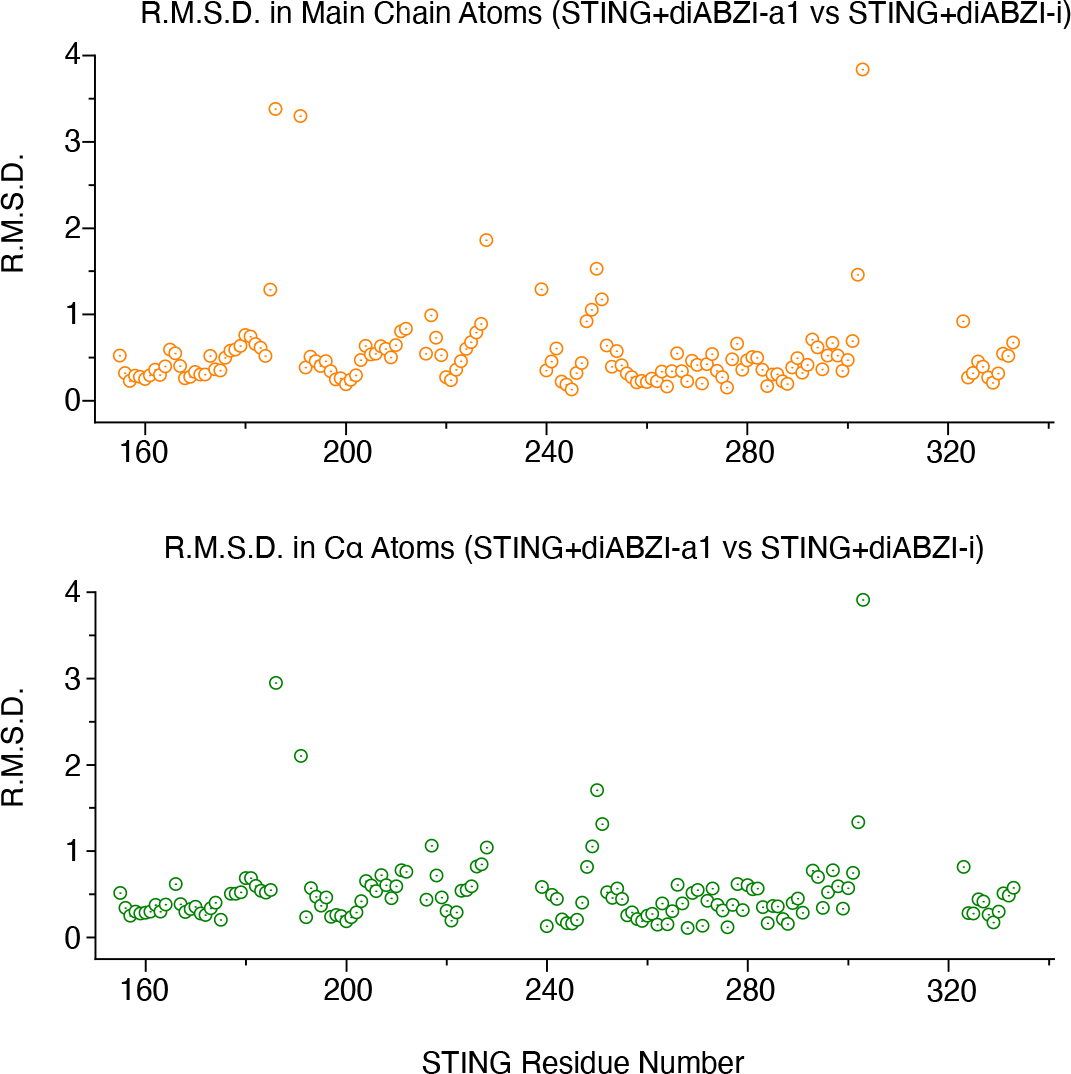
Root mean squared deviation (R.M.S.D.) between diABZI-i and diABZI- a1 of main chain atoms and C*α* atoms.

**Extended Fig. 2.**
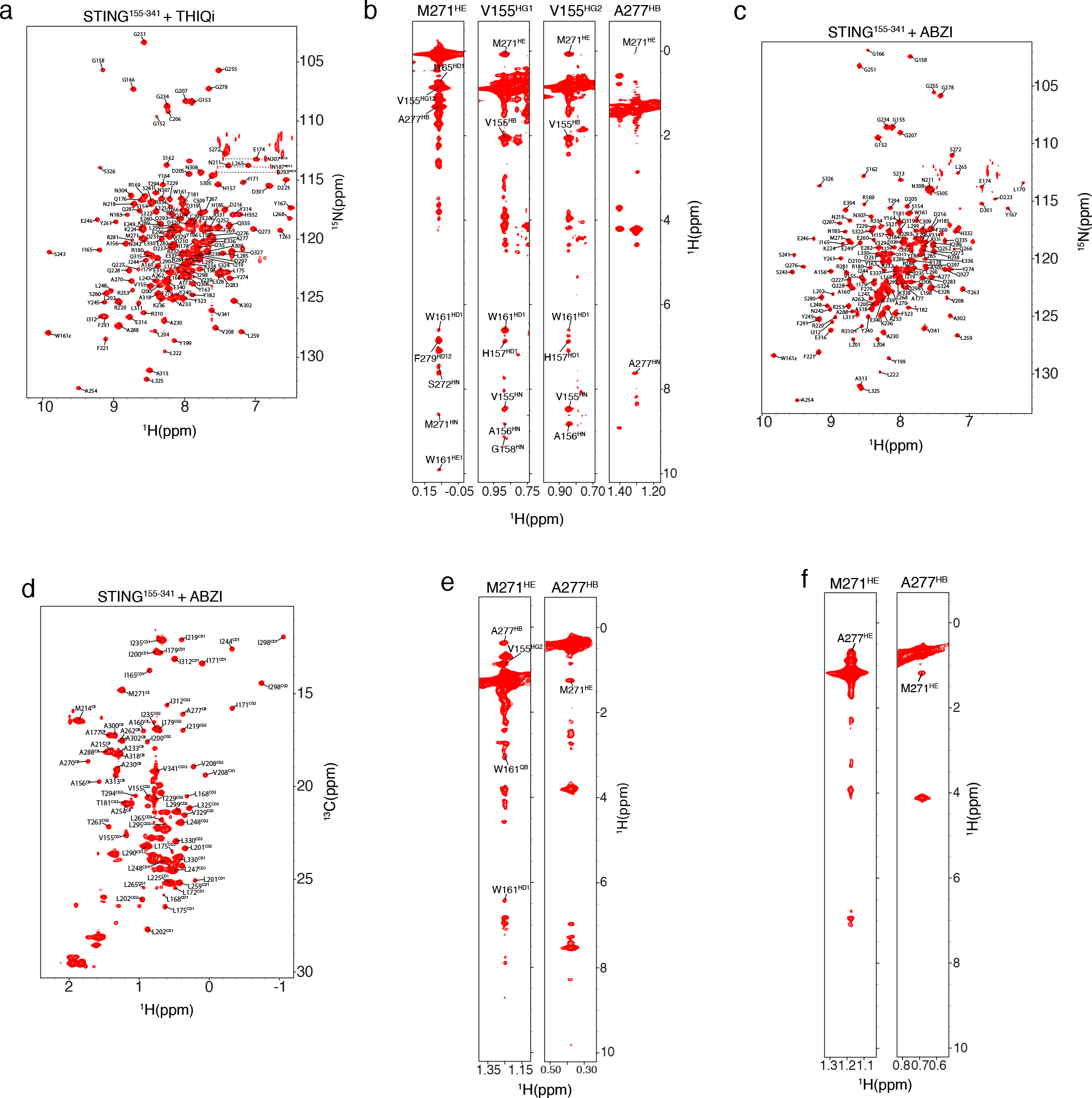
NMR spectra used for resonance assignments. (a) ^1^H-^15^N HSQC spectrum of STING^155-341^-THIQi complex annotated with backbone amide assignments. **(b)** Selected strips of ^13^C-edited NOESY-HSQC spectra of THIQi-bound STING^155-341^ highlighting the intramolecular NOEs between M271^CH3^ and other atoms. **(c)** ^1^H-^15^N HSQC and **(d)** ^1^H-^13^C HSQC of ABZI-bound STING^155-341^ annotated with assignments. **(e)** Selected strips of ^13^C-edited NOESY-HSQC spectra of ABZI-bound STING^155-341^ showing NOEs between M271^CH3^ and other atoms. **(f)** Selected strips of ^13^C-edited NOESY-HSQC spectra of diABZI-a1-bound STING^155-341^ confirming the assignment of A277^CH3^.

**Extended Fig. 3.**
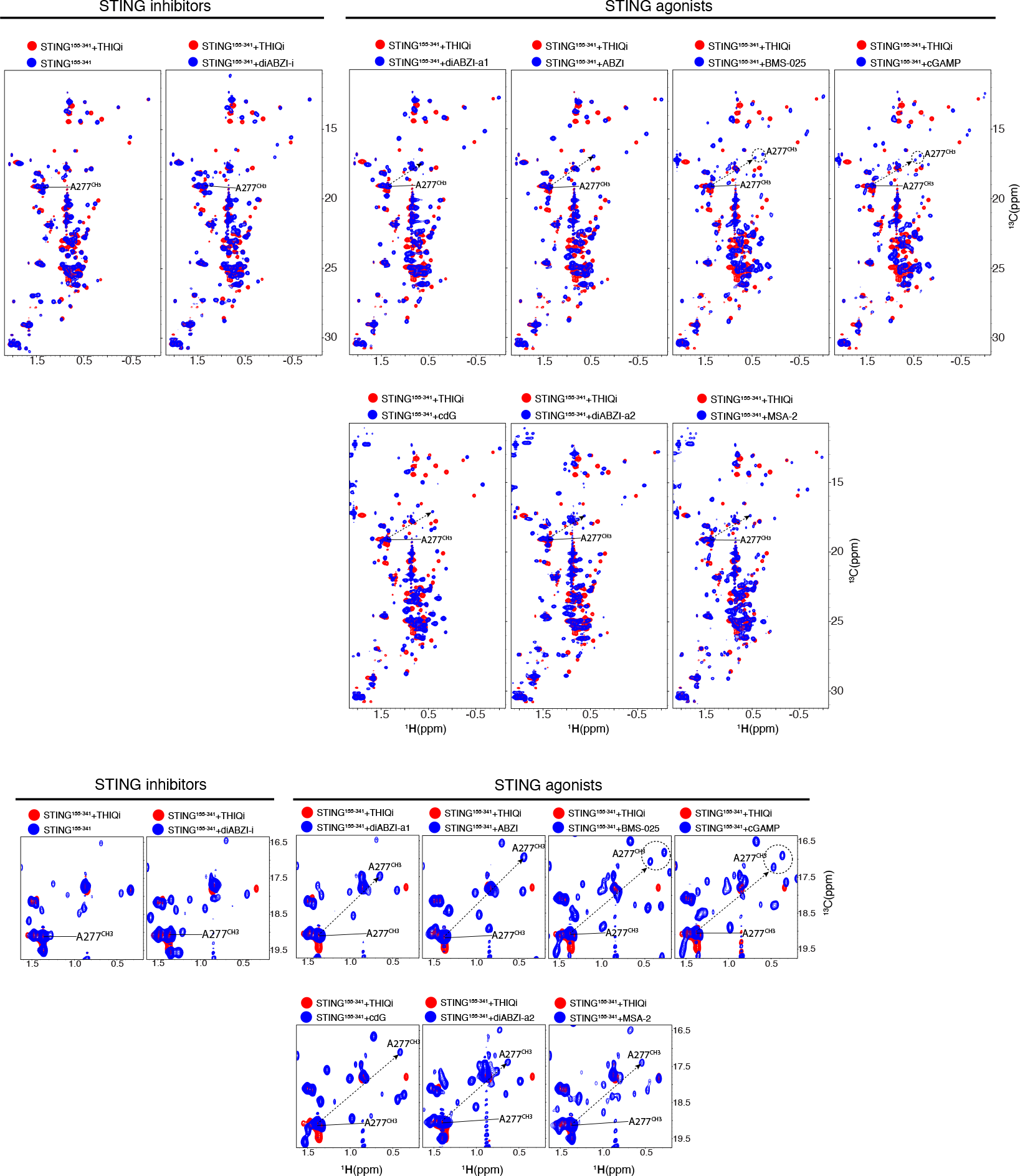
Overlay of ^1^H-^13^C HSQC of STING^155-341^ recorded in the absence and presence of various inhibitors and agonists.

**Extended Fig. 4.**
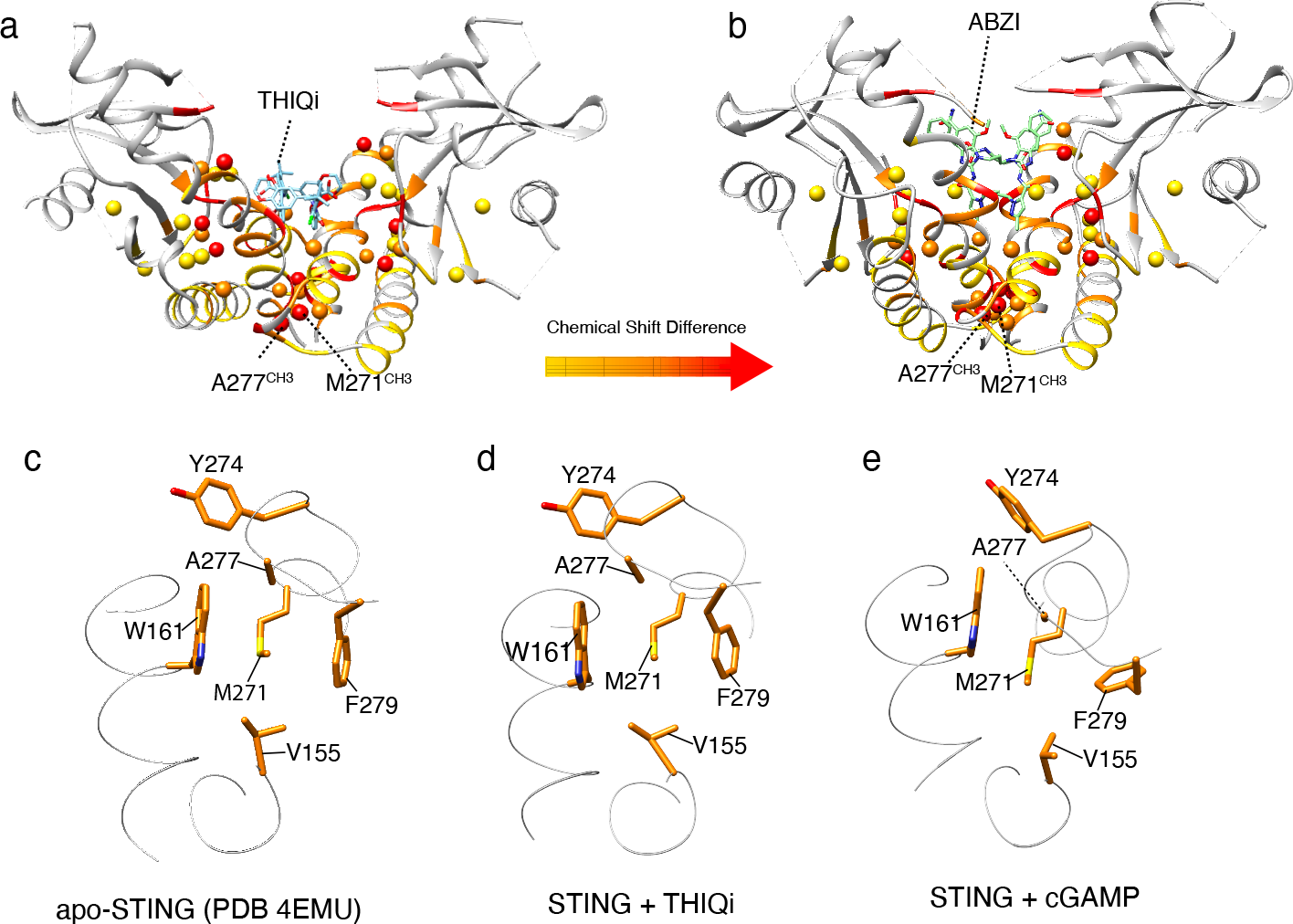
Structural basis for large chemical shift changes of M271^CH3^ and A277^CH3^ upon agonist binding. Chemical shift differences are mapped onto (a) STING^155-341^-THIQi and **(b)** STING^155-341^-ABZI structures. M271^CH3^ and A277^CH3^ exist underneath the binding pocket and are distant from the CDN binding pocket yet show significant chemical shifts differences, indicating different conformations between these two complexes. Hydrophobic interaction network in **(c)** apo-STING^155-341^, **(d)** THIQi-bound STING^155-341^, and **(e)** 2′,3′-cGAMP bound _STING_155-341.

**Extended Fig. 5.**
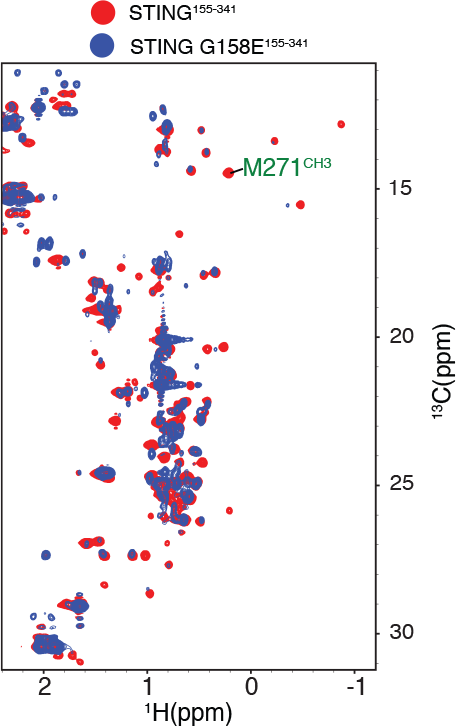
^1^H-^13^C HSQC spectrum comparison between STING^155-341^ (red) and STING^155-341^ G158E (blue).

## Notes

### Competing Interest Statement

The authors have declared no competing interest.

